# Systematic evaluation of transcriptomics-based deconvolution methods and references using thousands of clinical samples

**DOI:** 10.1101/2021.03.09.434660

**Authors:** Brian B. Nadel, Meritxell Oliva, Benjamin L. Shou, Keith Mitchell, Feiyang Ma, Dennis J. Montoya, Alice Mouton, Sarah Kim-Hellmuth, Barbara E. Stranger, Matteo Pellegrini, Serghei Mangul

## Abstract

Estimating cell type composition of blood and tissue samples is a biological challenge relevant in both laboratory studies and clinical care. In recent years, a number of computational tools have been developed to estimate cell type abundance using gene expression data. While these tools use a variety of approaches, they all leverage expression profiles from purified cell types to evaluate the cell type composition within samples. In this study, we compare ten deconvolution tools and evaluate their performance while using each of eleven separate reference profiles. Specifically, we have run deconvolution tools on over 4,000 samples with known cell type proportions, spanning both immune and stromal cell types. Twelve of these represent *in vitro* synthetic mixtures and 300 represent *in silico* synthetic mixtures prepared using single cell data. A final 3,728 clinical samples have been collected from the Framingham Cohort, for which cell populations have been quantified using electrical impedance cell counting. When tools are applied to the Framingham dataset, the tool EPIC produces the highest correlation while GEDIT produces the lowest error. The best tool for other datasets is varied, but CIBERSORT and GEDIT most consistently produce accurate results. In terms of reference choice, we find that the Human Primary Cell Atlas (HPCA) and references published by the EPIC authors produce accurate results for the largest number of tools and datasets. When applying deconvolution to blood samples, the leukocyte reference matrix LM22 is also a suitable choice, usually (but not always) outperforming HPCA and EPIC. Running time varies substantially across tools. For as many as 5052 samples, SaVanT and dtangle reliably finish in under one minute, while slower tools may require up to two hours. However, when using custom references, CIBERSORT can run very slowly, taking over 24 hours to complete for large datasets. We conclude that combining the best tools with optimal reference datasets can provide significant gains in accuracy when carrying out deconvolution tasks.

## Introduction

Biological tissues are rarely homogeneous, and are instead typically composed of a variety of distinct cell types. The relative abundance of these cell types is fundamental to tissue biology and function, and therefore of frequent interest to the biomedical community. In medical settings, knowledge of cell type populations can provide insight into the nature of a wide range of diseases and, in some cases, inform treatment. In cancer, for instance, the abundance of certain T cells correlates strongly with survival, as well as the efficacy of immunotherapy treatment^1–3^. In laboratory settings, researchers frequently observe gene expression changes that are difficult to interpret without knowledge of cell type composition. Such patterns may be the result of changes of cell type abundance, rather than altered expression in any particular cell type. Cell type deconvolution methods enable researchers to distinguish between these two cases and extract further insights from their experiments.

Existing molecular techniques of cell type quantification can be difficult to apply to large scale studies or to certain cell types. Fluorescence-Activated Cell Sorting (FACS) is often considered a gold standard method. However, FACS is typically based on a limited number of markers that are selected beforehand, sometimes limiting the cell types that can be quantified. Moreover, standard FACS cannot quantify cell types with unusual morphologies, such as neurons, myocytes, and adipocytes. Single cell RNA-seq methods (scRNA-seq) are becoming increasingly popular, but their cost remains high^4^. Moreover, current scRNA-seq methods capture only a small fraction of cells present in a tissue, and the observed cells may not represent a random sample^5^. Both scRNA-seq and FACS require cells to be dissociated into a single cell suspension before processing ^6^. During this process, some cells may be lysed before they are observed, while others remain aggregated and are less likely to be detected. Consequently, subtle alterations during the cell dissociation step can produce dramatic differences in observed cell type fractions^7^.

To overcome these limitations, a number of expression-based methods have been developed that aim to serve the biomedical community’s need for accurate estimation of cell type abundances from gene expression data ^8–15^. These tools utilize either RNA-seq or microarray expression data to digitally deconstruct tissue samples, a process known as cell type deconvolution. However, it is often unclear to the user which tool will best suit their needs^16^. Evaluations performed as part of tool publications are often limited in scope, assessing accuracy for only a limited number of datasets, platforms, and tissue types.

Attempts to benchmark these tools have largely been limited to simulated data, which fails to capture the true complexity of tissue samples in living organisms ^16,17^. Studies that do incorporate clinical data often rely on small numbers of samples or on samples of unknown cell type content (rather than evaluated by reliable molecular methods). Conclusions derived from these limited datasets can be misleading and incomplete, and we currently lack a systematic comparison of deconvolution methods evaluated on high-quality data. As such, researchers are left with little guidance as to which deconvolution tool is most suitable for their needs.

In addition, reference data is a requirement for many tools, and existing benchmarking studies do not address the relationship between choice of reference and prediction quality. Here, we explore this relationship by running tools using a variety of reference datasets and reporting performance in each case.

In this study, we compare ten deconvolution tools and evaluate their performance across eleven separate reference profiles. We have tested all these tools using multiple references, and have produced predictions for 3,728 clinical blood samples, 300 simulated blood and stromal samples, and 12 blood samples mixed *in vitro*. The cell type composition of the clinical blood samples has been evaluated via an impedance-based electronic cell counter, a gold standard for high-throughput cell type quantification in blood. In addition, we have thoroughly evaluated the effect of reference choice on the accuracy of deconvolution predictions. Overall, we utilize over 3,500 clinical samples from the Framingham cohort study ^18–20^ to evaluate the performance of deconvolution methods in the most powerful, complete and unbiased manner to-date.

## Results

### Deconvolution tools selected for benchmarking

We have selected ten commonly used deconvolution tools, which we benchmarked on our datasets in order to evaluate their ability to accurately predict cell type content of tissue samples. The tools included in this study are CIBERSORT (normal and absolute mode)^10^, the DCQ algorithm^9^, DeconRNASeq^8^, dtangle^13^, EPIC^15^, GEDIT, MCP-Counter^11^, quaNTiseq^14^, SaVanT^22^, and xCell^12^ (Table 2). Some of these tools do not explicitly predict fractions, but rather ‘enrichment scores’. These scores may be calculated based on the sum (or log-sum) of marker gene expression, and attempt to quantify the prominence of cell type signatures in each sample.

While scores for a particular signature can be compared across samples, comparisons across cell types may not be valid and are not recommended. For example, a higher score for B cells compared to NK cells does not imply a greater prevalence of B cells. Other tools perform deconvolution, utilizing such tools as basic linear regression (e.g. DeconRNASeq), modified linear regression (e.g. dtangle, quaNTiseq) and support vector regression (CIBERSORT). Outputs for these tools can be interpreted as fractions, allowing comparisons across cell types.

### Gold standard datasets

In this study, we use both simulated and experimental datasets to evaluate the accuracy of existing deconvolution tools (Supplementary Table 1). Cell type fractions of these datasets have been evaluated by trusted molecular techniques able to deliver highly accurate cell type proportions. The samples consist of 300 synthetic mixtures prepared *in silico* using single cell data, 12 mixtures prepared *in vitro* and sequenced using microarray, and 3,728 clinical samples.

The 300 pseudo-bulk synthetic mixtures were prepared using scRNA-seq data. For each mixture, individual cells were randomly selected and their expression profiles summed: 200 of these represent simulated PBMC mixtures containing five common PBMC cell types (B, CD4 T, CD8 T, natural killer and monocytes). Exactly 100 of these were created using data obtained from a previous study^21^ and 100 using data from 10x Genomics (https://www.10xgenomics.com/resources/datasets/). Lastly, a third set of 100 mixtures contained stromal cell types as well (B, CD4 T, CD8 T, macrophage, mast, endothelial and fibroblast cells), also using data from 10x Genomics. Following clustering, PBMC cell type assignment was performed using four to six marker genes for each cell type (Supplementary Figure 1). Cell type assignment of the stromal dataset was performed by 10x Genomics. In each case, 1000 cells were selected in total, their expression values summed, and the cell type ratios noted.

In addition, 12 *in vitro* mixtures were prepared by physical titration of purified cell types in known proportions. Six immune cell types were used to produce these mixtures (B, CD4 T, CD8 T, monocytes, natural killer, neutrophils), which were combined in varying proportions (Supplementary Figure 2). The mixtures were then profiled using an Illumina HT12 BeadChip microarray. Lastly, the Framingham Cohort data ^20^ is collected from the blood of healthy individuals and profiled on Affymetrix Human Exon Array ST 1.0 arrays. Gold standard cell type fractions were obtained using electrical impedance.

### Optimal reference choice varies across deconvolution tools

In addition to expression data from heterogeneous tissue samples, most deconvolution tools require reference data containing expression profiles of pure cell types. These data can be in the form of an expression matrix, a list of signature genes, or both. It has been shown that the choice of reference can have a significant impact on the quality of results ^23^, and as part of this study we explore the effect of reference choice on tool performance.

As reference data is a requirement for most tools included in this study, we have carefully curated an extensive list of reference datasets (Supplementary Table 2). These references include data from a variety of sources, including scRNA-seq, bulk RNA-seq, and multiarray platforms. Each reference contains a different set of cells, with some including stromal cells and others only immune. Each reference also contains a varying number of genes; ImmunoStates^24^ and LM22^10^ have been curated to contain only a short list of signature genes, whereas other matrices contain the larger set of genes measured by their respective platforms. In this study, there are eight deconvolution tools that accept custom references and we provide ten reference datasets to each. The reference profiles used derive from a variety of sources and platforms, including LM22^10^, ImmunoStates^24^, 10x Genomics^25^, EPIC^15^, BLUEPRINT^26^, and the Human Primary Cell Atlas^27^. In order to evaluate the effect of reference choice on prediction quality, we systematically evaluate each tool using each reference source. For every set of results, we calculate the Pearson correlation between true cell type fractions and tool output, and treat these correlations as our metric of performance. For tools that predict fractions, we also compare average absolute error (Supplementary Figure 3). We find that the choice of reference has a substantial impact on quality of results across all datasets (Figure 1). Some tools produce accurate results when the optimal reference is selected, but high error and even negative correlations when an improper reference is used. This is particularly evident when applied to the Framingham dataset, where several references produce poor-quality predictions when used with any tool (10x Immune, BLUEPRINT, EPIC-TIC, HPCA-Stromal). While no reference performs best for all tools or all mixtures, LM22 may be the most suitable for predicting hematopoietic cell types. EPIC and HPCA references also perform well in many cases, and allow the user to predict fractions of both stromal and blood cell types.

**Figure 1.**
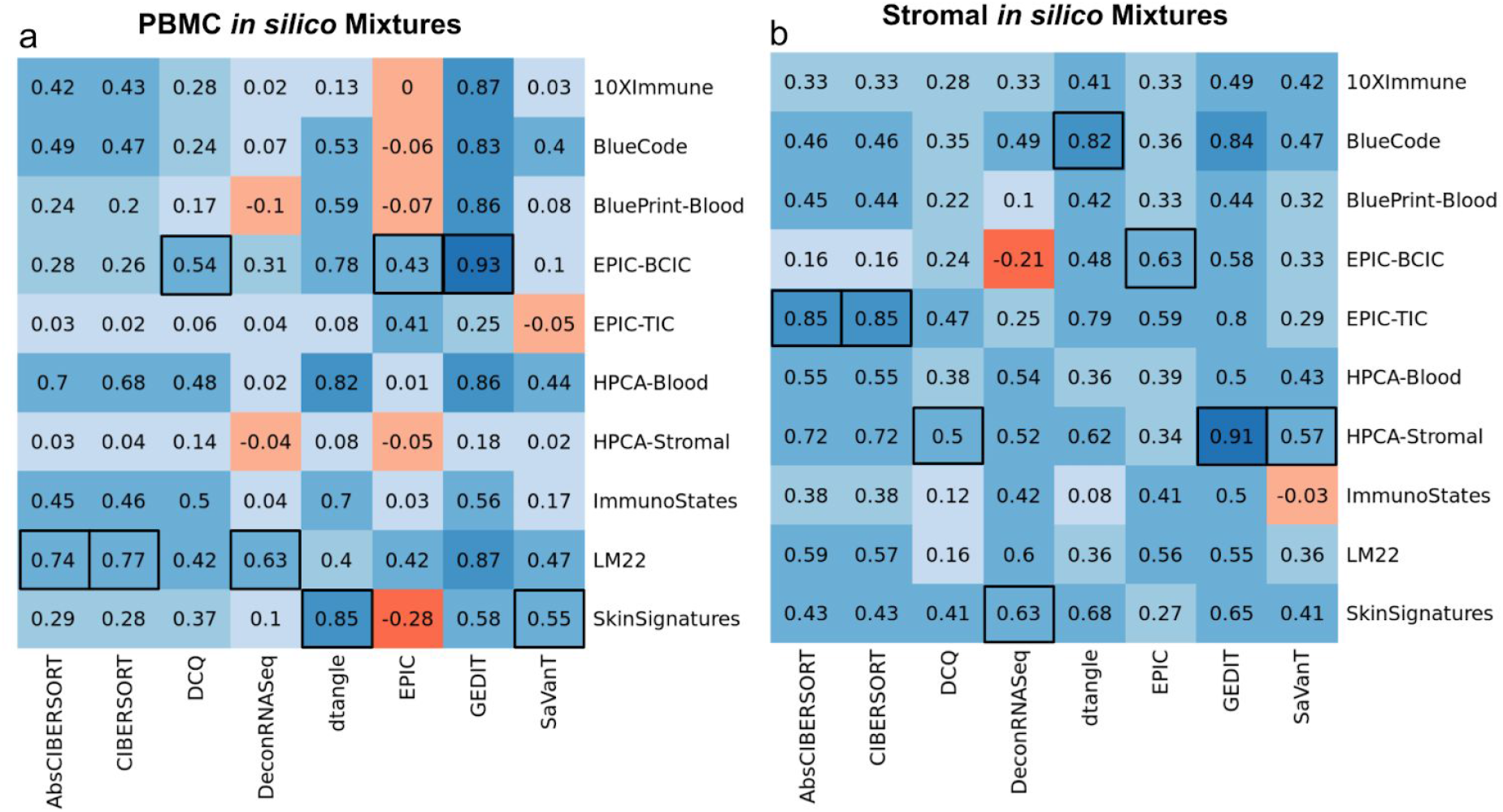
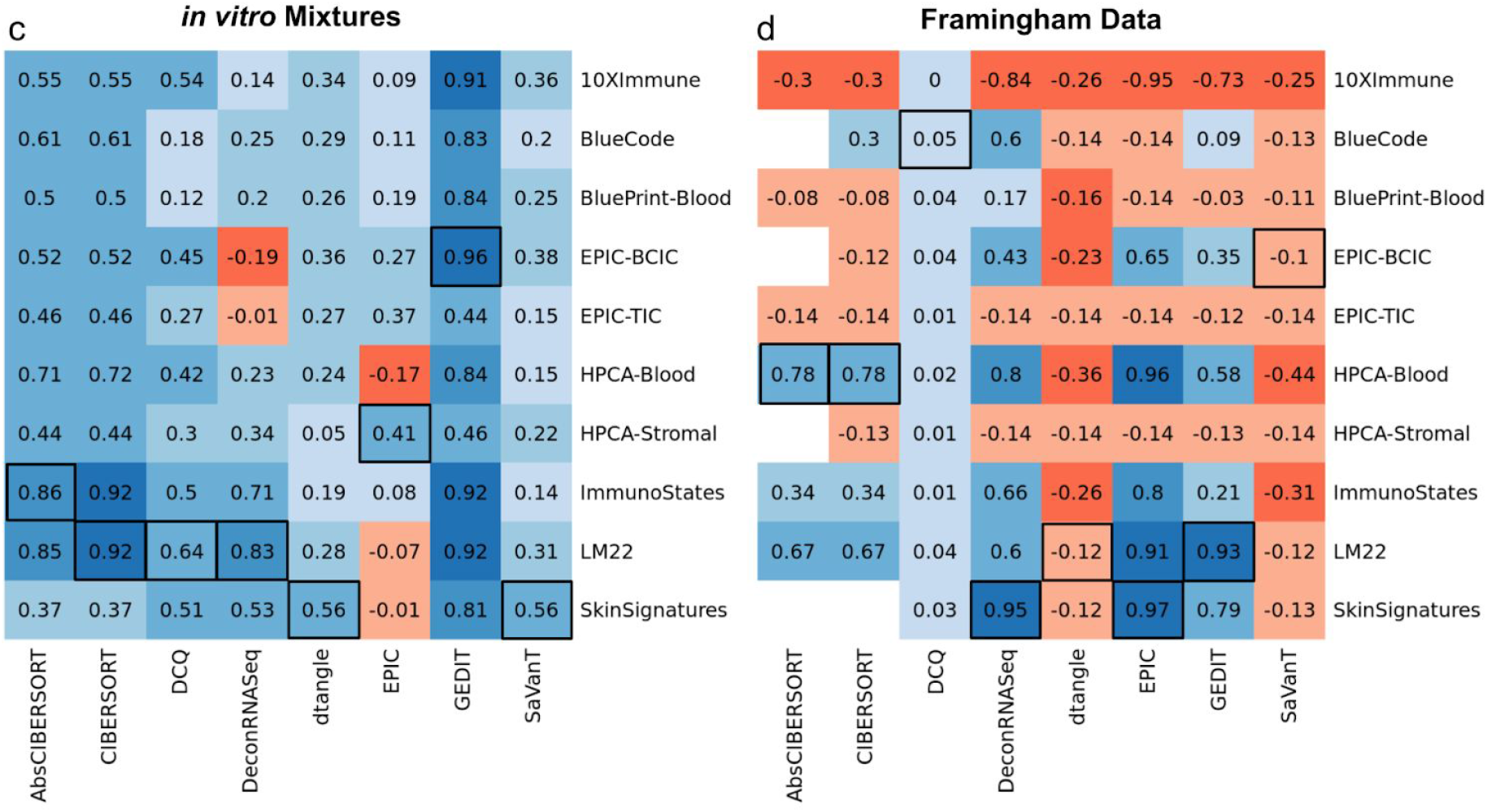
Determining the optimal reference choice for each tool that supports external references. Shown here are Pearson correlations between true cell fractions and tool output for each combination of tool and reference matrix. Results are shown for PBMC and Stromal *in silico* simulated mixtures (a and b), for *in vitro* mixtures of immune cells (c), and clinical samples from the Framingham cohort dataset (d). Tools not able to deliver results within 48 hours were excluded and are not reported here. Highest correlation for each tool is shown in black boxes.

### CIBERSORT and GEDIT produce consistently high correlations between estimated and true cell type fractions

Next, we compare the performance of deconvolution tools when the optimal reference profile is used. We utilize two metrics to evaluate the accuracy of predicted fractions compared to actual fractions: correlation and absolute error. As performed in the previous section, we compute the Pearson correlation between true cell type fractions and tool output. These outputs can represent either estimated cell type fractions or enrichment scores.

Based on correlation analysis, CIBERSORT and GEDIT produce the most accurate results across all datasets, though both are outperformed by DeconRNASeq and EPIC when applied to the Framingham dataset. For each mixture tested, multiple tools produce highly accurate results (correlation greater than .9), though no single tool is able to maintain the highest performance across all datasets.

### CIBERSORT and GEDIT produce the lowest absolute errors across all datasets

We compare the ability of each tool to specifically predict cell type fractions, rather than scores. In order to produce predicted fractions using tools that are not designed to do so, we perform a simple transformation to convert the output into predicted fractions. For each sample, we divide each cell-type score by the sum of scores for all cell types. Therefore, the resulting sum is 1.0, and can be treated as fractions. Though this approach falls outside of the intended usage of certain tools (specifically, DCQ, MCP-Counter, and SaVaNT), it is capable of producing accurate results in some circumstances (Figure 3).

**Figure 2.**
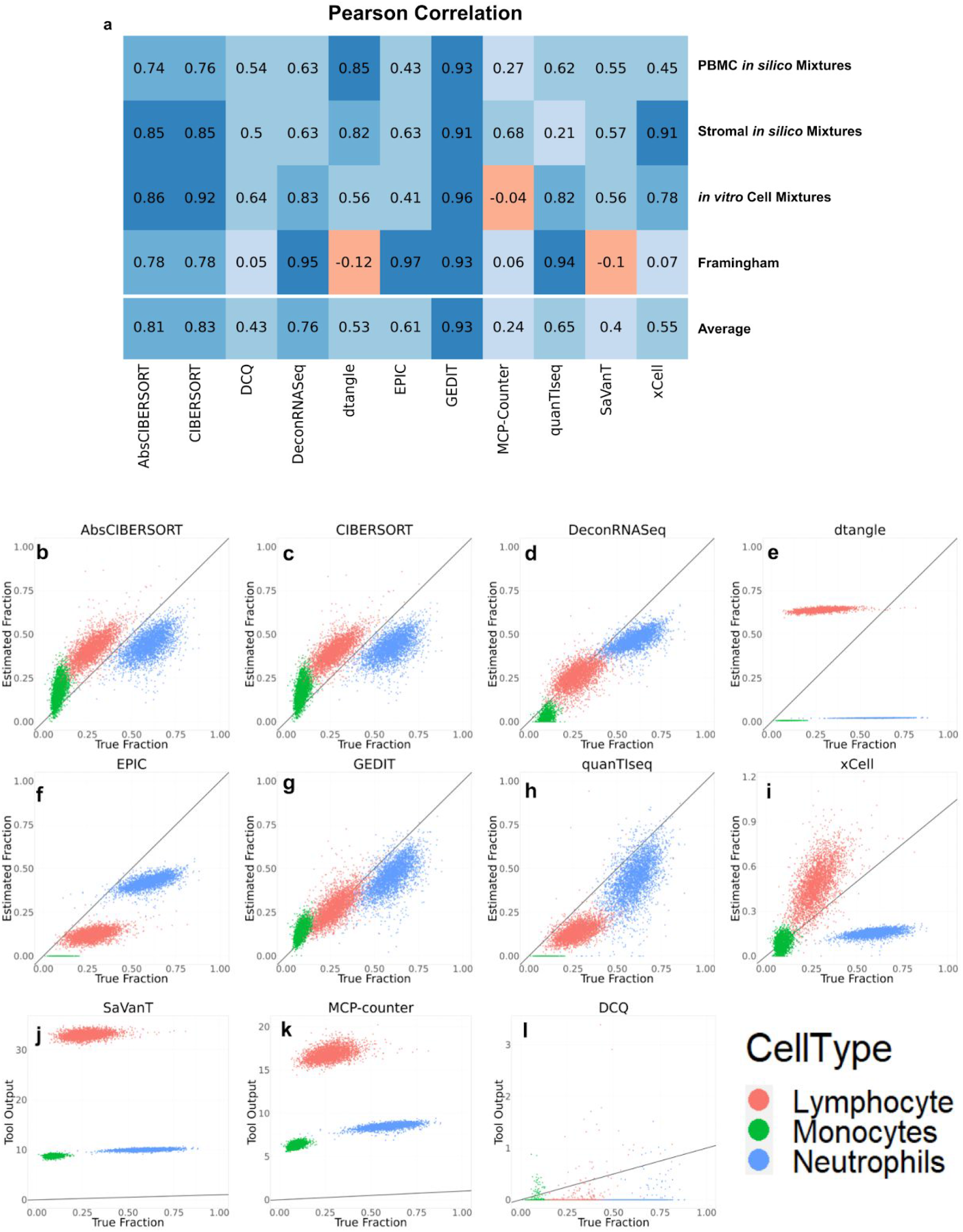
Comparisons between tool outputs and true cell type fractions, as evaluated by gold standard techniques. a) Pearson correlations between tool output and true fractions for each combination of tool and dataset; here, the optimal reference is used for each dataset (see Figure 1). b-l) Scatter plots visualizing the output of each deconvolution tool (y-axis) versus cell type fractions as evaluated by automated cell counting (x-axis). Three cell types were evaluated, with lymphocytes, monocytes, and neutrophils shown in red, green and blue, respectively. The y=x line is shown in black. For tools that accept custom reference data, the reference data that resulted in the highest correlation is shown here (see Figure 1d). Equivalent graphs for the other three datasets are included in supplementary materials (Supplementary Figures 4-6).

**Figure 3.**
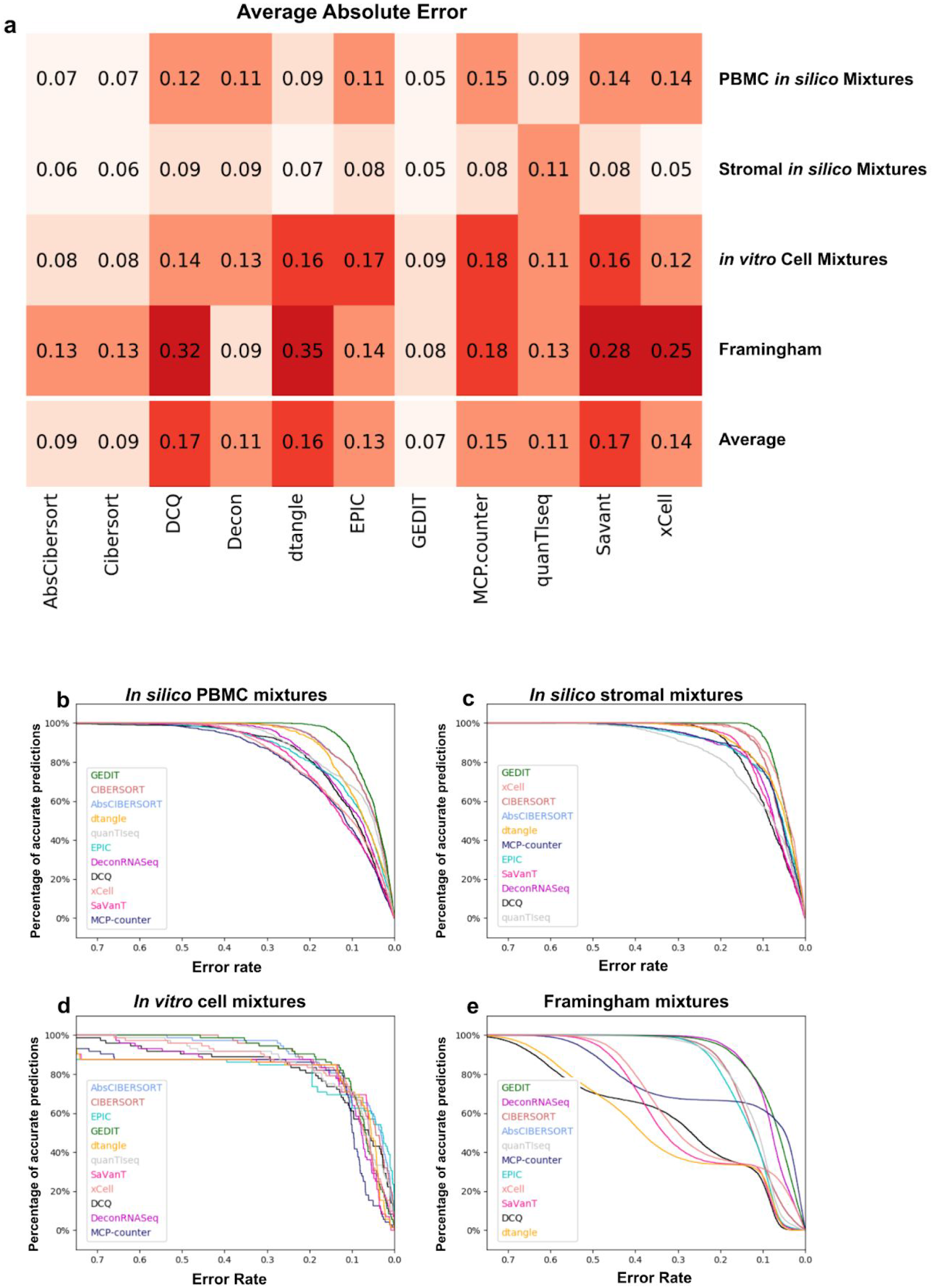
Error values of predictions for each tool and dataset. a) Absolute error values, averaged across all predictions, for each tool and dataset. b-e) distribution of absolute error for all predictions in each dataset. Predictions are considered accurate (y-axis) if error is less than allowed error rate (x-axis). Legend is sorted by decreasing area under the curve (Supplementary Table 3). For each combination of tool and dataset, the reference producing the highest correlation value was used (Figure 1).

**Figure 4.**
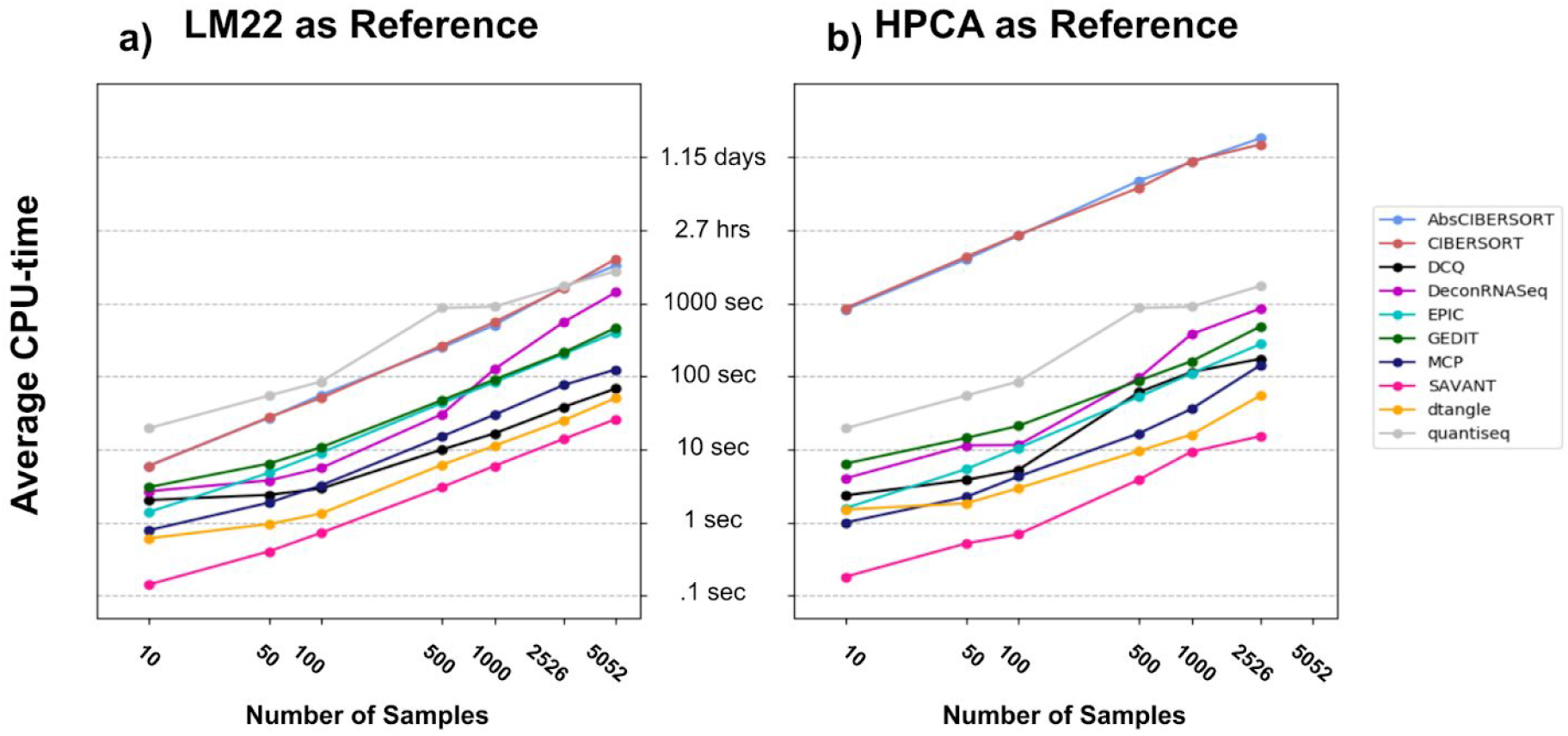
Time requirements for each tool for varying numbers of samples. Subsamples of varying size were randomly selected from the Framingham dataset. Each tool was run twice using LM22 (a) and the Human Primary Cell Atlas (b) as reference data; the larger HPCA produces longer runtimes in most cases. Each point represents the average of 20, 10, 10, 5, 5, 2 runs for input sizes of 10, 50, 100, 500, 1000, 2526, respectively. A single sample of size 5052 was also run for all tools using LM22 as the reference input. In both cases, the default settings for MCP-Counter and quanTIseq were used, as they do not support external references.

The error of predicted fractions varies greatly depending on the exact combination of tool and mixture. CIBERSORT and GEDIT are able to maintain high accuracy across the majority of datasets. For all datasets, the ‘absolute’ mode of CIBERSORT produces slightly higher error than the default mode. This is likely because the default CIBERSORT is designed to predict fractions, whereas the absolute version is not. xCell performs best on the *in silico* simulated mixtures, but produces high error for some cell types in the *in vitro* mixture (e.g. B Cells).

For all tools, we observe some of the highest errors when applied to the Framingham data. This is likely due to complexities of biological samples, including varying sub-states for many cell types that are not adequately reproduced in simulated data (whether *in vitro* or *in silico*).

### Runtime varies substantially depending on tools and references

We evaluate the scalability of deconvolution by varying the size of inputs and recording the CPU time required by each tool. Specifically, we randomly subsample the Framingham Cohort data into batches varying in size from 10 to 5052 samples. We then record CPU time required to run each tool as a function of input size. We exclude xCell from this analysis, since it does not support an easily accessible command line interface.

Further, we explore the effect of reference choice on resource requirements (for the tools that support custom references). Specifically, two separate references are applied and runtime recorded in each case; these references are the larger HPCA reference containing 19,715 genes, and the smaller LM22 reference containing only 547 genes. We find that the size of the reference matrix has a substantial effect on the running time of certain tools, in particular CIBERSORT (both absolute and default moudes). When the smaller reference is used all tasks complete relatively quickly, with the slowest run taking 1.1 hours. However, when the larger HPCA reference is used, runtimes for some tools (specifically, both modes of CIBERSORT) can reach over 24 hours for large numbers of samples.

## Discussion

Transcriptomics-based cell type deconvolution is an increasingly popular approach for estimating the cell type composition of heterogeneous samples. However, current tools and reference profiles are numerous, and it is important that researchers have a clear way of determining the best choices for their needs. Here, we perform a comprehensive benchmarking study to systematically evaluate the performance of various computational deconvolution tools across 4,040 transcriptomics samples using accurate molecularly defined gold standard. We explore the effects of reference profiles and demonstrate that choice of reference can have a substantial impact on performance of the tool. Some tools demonstrate increased sensitivity to the choice of reference compared to other tools. For example, dtangle can produce accurate predictions when the correct reference is used, but performs poorly with other reference choices.

Moreover, the best choice of reference varies between tools, even when applied to the same mixture. Utilizing GEDIT combined with the EPIC ‘Blood Circulating Immune Cells’ reference produces the most accurate results for the *in vitro* mixtures dataset. However, this same reference produces highly inaccurate predictions when combined with the DeconRNASeq tool. The best results for DeconRNAseq are obtained when the LM22 reference is used. Several other tools produce their best results when utilizing the LM22 or Skin Signatures references. LM22 is perhaps the most robust reference, as it is the least likely to produce poor correlations or high error between predicted and actual cell type proportions. When applying deconvolution to stromal data, the EPIC Tumor Infiltrating Cells reference yields the best average performance across tools. However, the best overall results come from GEDIT while using the HPCA-Stromal reference.

Most tools return results within 30 minutes, even on datasets larger than 5,000 samples.. However, DeconRNASeq and CIBERSORT (both modes) have potential for long runtimes, potentially taking hours or even days to return results. For reference based tools, runtimes can often be reduced by supplying reference matrices containing a small number of signature genes, rather than the complete transcriptome.

While no single tool produces the best results across all types of datasets, CIBERSORT (both modes), DeconRNASeq, and GEDIT are able to produce reliable results (average error less than 0.15, correlation greater than 0.6) across all four datasets. However, proper reference selection is critical, with EPIC-BCIC, HPCA-blood and LM22 returning accurate results for blood samples, and EPIC-TCIC or HPCA-stromal performing well for samples containing stromal cells. For the Framingham dataset, the largest dataset in the study, DeconRNASeq, EPIC, quaNTiseq and GEDIT all produce very high correlations between predicted and actual fractions, with DeconRNASeq and GEDIT producing the lowest errors.

Taking all datasets into account, GEDIT and CIBERSORT are able to acheive high accuracy across all datasets. By the metrics used in this study, the default mode of CIBERSORT outperforms the “absolute” mode by a small margin. Interestingly, the relatively simple linear regression algorithm of DeconRNASeq outperforms several more complex methods, provided that the proper reference is used. Remarkably, several tools perform poorly on the Framingham dataset, producing high errors and even negative correlations.

Several regression based tools (CIBERSORT, DeconRNASeq, GEDIT) and reference data sources (LM22, EPIC, HPCA) produce reliable accurate results for all datasets. For many tools, however, prediction quality varies dramatically depending on both the provided dataset and reference source. As such, researchers applying deconvolution to novel datasets (and especially novel cell types) may wish to run deconvolution using multiple tools and/or references. By examining the consistency of results across multiple conditions, one can differentiate between real biological patterns and technical artefacts.

## Methods

### PBMC and Stromal Single Cell Mixtures

We obtained 2 datasets from 10x Genomics (https://www.10xgenomics.com/resources/datasets/) and 1 from GEO ^21^.

The PBMC mixtures were created in two batches, each producing 100 mixtures. For the first set, we used 1000 cells for each sorted cell type (monocytes, CD8 and CD4 T cells, B cells, natural killer cells). For each cell type, we randomly selected 1-1000 cells, then we sum the expression of all the selected cells to create a synthetic mixture. The process was repeated 100 times, thus 100 mixtures were created. For PBMC2, we firsted clustered the cells and identified the cell types for the dataset. Then we used the same five cell types in PBMC2 and created the 100 mixtures the same way as we did from PBMC1. For stromal cells, we created 100 mixtures the same way as we did for PBMC2 except for that we included two additional cell types not typically found in blood (Fibroblasts and Mast cells), and exclude monocytes and natural killer cells. For each mixture, the true fraction for each cell type is calculated as the number of cells of that cell type selected, divided by the total number of cells across all cell types.

### Framingham Data

The Framingham Heart Study (FHS) is a population-based study, predominantly of European ancestry, consisting of an ongoing series of primarily family-based cohorts first developed in 1948 and based in Framingham, MA, USA; it comprises the Original ^18^, Offspring ^19^, and Third Generation ^20^ cohorts. FHS gene expression, blood cell counts, subject and sample metadata was obtained from dbGap (phs000007). Gene expression data was processed (filtered and normalized) as in ^28^, resulting in 5,058 available FHS samples, of which 3,728 had available blood cell counts. Blood cell counts were obtained through a Complete Blood Count using the Coulter HmX Hematology Analyzer (Beckman Coulter, Inc.) ^29,30^.

The “gold standard” is cell counts and cell percent. The cell counting was performed on a Beckman Coulter HmX hematology analyzer. The following metrics from whole blood were obtained - HbA1c, basophil count and percent, eosinophil count and percent, hematocrit, hemoglobin, lymphocyte count and percent, MCH, MCHC, MCV, monocyte count and percent, MPV, neutrophil count and percent, platelet count, RBC, RDW, and WBC. Further information is available here: https://www.ncbi.nlm.nih.gov/projects/gap/cgi-bin/GetPdf.cgi?id=phd004086.1.

### Selected tools

We have selected available deconvolution tools able to infer the relative abundances of immune cell types based on the gene expression profiles. In total, we have identified 9 deconvolution tools which either predict cell type fractions or produce scores. xCell was run using the online interface at https://xcell.ucsf.edu/, but all other tools were installed on the Hoffman2 Cluster at UCLA. Each deconvolution task was provided 16Gb of RAM and 48 hours of runtime.

#### CIBERSORT

We requested the CIBERSORT code from the author’s website https://cibersort.stanford.edu/download.php and have installed CIBERSORT version 1.04 on the UCLA Hoffman2 cluster (R version 3.6.0). We ran all deconvolution tasks using both the default CIBERSORT mode, and the absolute mode, and reported both results. As the statistical outputs of CIBERSORT (e.g. p-values) are not considered in our analysis, we ran with 0 permutations to reduce resource usage. Quantile normalization was used for the Cell Mixtures and Framingham datasets, but disabled for the PBMC and Stromal datasets; this follows the author recommendations regarding application to microarray and RNA-seq data, respectively. Several tasks involving the large Framingham data failed to complete due to high resource usage. In these cases, we split this 5053 sample dataset into three smaller datasets, each with 1684 samples, and ran three deconvolution tasks separately.

#### DCQ

DCQ was installed as part of the ComICS package in R. For each reference matrix used, we designated all genes present in that matrix as marker genes.

#### DeconRNASeq

The DeconRNASeq R package was installed from Bioconductor, and run using default settings.

#### dtangle

The dtangle R package (version 2.0.9) was obtained from CRAN and installed on the Hoffman2. As part of the wrapper to run the function, genes not shared between the mixture and reference matrices were excluded (otherwise this caused a crash). All observed values X in both matrices were then transformed to log2(1+x), as dtangle takes log transformed data as input. The dtangle() function was used with default settings.

#### EPIC

We downloaded EPIC from the author’s GitHub repository (https://github.com/GfellerLab/EPIC) and installed it on the Hoffman2 Cluster. The tool provides two built in reference datasets (Tumor Infiltrating Cells and Blood Circulating Immune Cells). When running the tool using these datasets, we use the default mode that utilizes additional data regarding reference profile variability and amount of mRNA per cell. For the other eight reference datasets used in this study, these additional data are not available and thus were not included as inputs. The cell fractions outputs are taken as cell type estimates.

#### GEDIT

GEDIT version 1.6 was obtained from the GitHub repository https://github.com/BNadel/GEDIT. Necessary packages were installed, specifically random, numpy, glmnet, RColorBrewer, and gplots. It was run using default settings.

#### Microenvironment Cell Populations-counter (MCPcounter)

The MCP-Counter code was obtained from the github (https://github.com/ebecht/MCPcounter) and installed on Hoffman2 along with its dependencies devtools and curl. The MCPcounter.estimate() function was used to produce predictions. “HUGO_symbols’’ was designated, and otherwise default settings were used. We contacted the authors regarding use of external references, but this appears to require direct modification of the MCP-Counter code, and was therefore not performed in this study.

#### quanTIseq

The quanTIseq code was obtained from the GitHub https://github.com/icbi-lab/quanTIseq and run on the Hoffman2 Cluster; installation via docker or singularity both failed on the cluster. Specifically, the quantiseq_pipeline.sh script was called using the command of the form “./quanTIseq_pipeline.sh --inputfile=$MIXTUREFILE --outputdir=$OUTPUTFILE --pipelinestart=decon”.

#### SaVanT

The code for SaVanT was obtained from it’s authors and run using 50 signature genes per cell type.

#### xCell

xCell was run using the online tool found at http://xcell.ucsf.edu/. The bulk expression data is submitted under “upload gene expression data” and the default gene signatures were used (xCell, n=64). The RNASeq option was selected for PBMC and Stromal datasets but not for CellMixtures or Framingham data.

#### Reference Data

Reference data was obtained from a variety of sources, as described in Supplementary Table 2. The HPCA reference matrix contained a wide variety of cells that were not present in any of our mixtures. As such, we subsetted this reference matrix in order to produce two versions more appropriate for the data used in the present study. These two versions contain 7 blood cell types (B, CD14+ monocytes, CD16+ monocytes, NK, neutrophils, CD4+ T and CD8+ T) and 6 blood and stromal cell types (B, CD4+ T, CD8+ T, endothelial, fibroblast, macrophage), respectively.

#### Cell Type Matching

Depending on the exact tool and reference matrix used, prediction outputs could be labelled as any one of 219 cell types and subtypes (Supplementary Table 4). For each output, we either match the output with a cell type quantified in the mixture, or note that it does not match any. In some cases this matching is trivial (e.g. B_cells, B-Cells, and BCells are all noted as “B Cells”), and in other cases the outputs represent cell subtypes (e.g. “naive B-Cells” and “memory B Cells” were also noted as “B Cells”). The final output for each cell type in the mixtures was calculated as the sum of outputs matched with that cell type; thus predictions for a general cell type are computed as the sum of the subtypes for that cell. The table of output interpretation is included as a supplementary file.

**Table 1.**
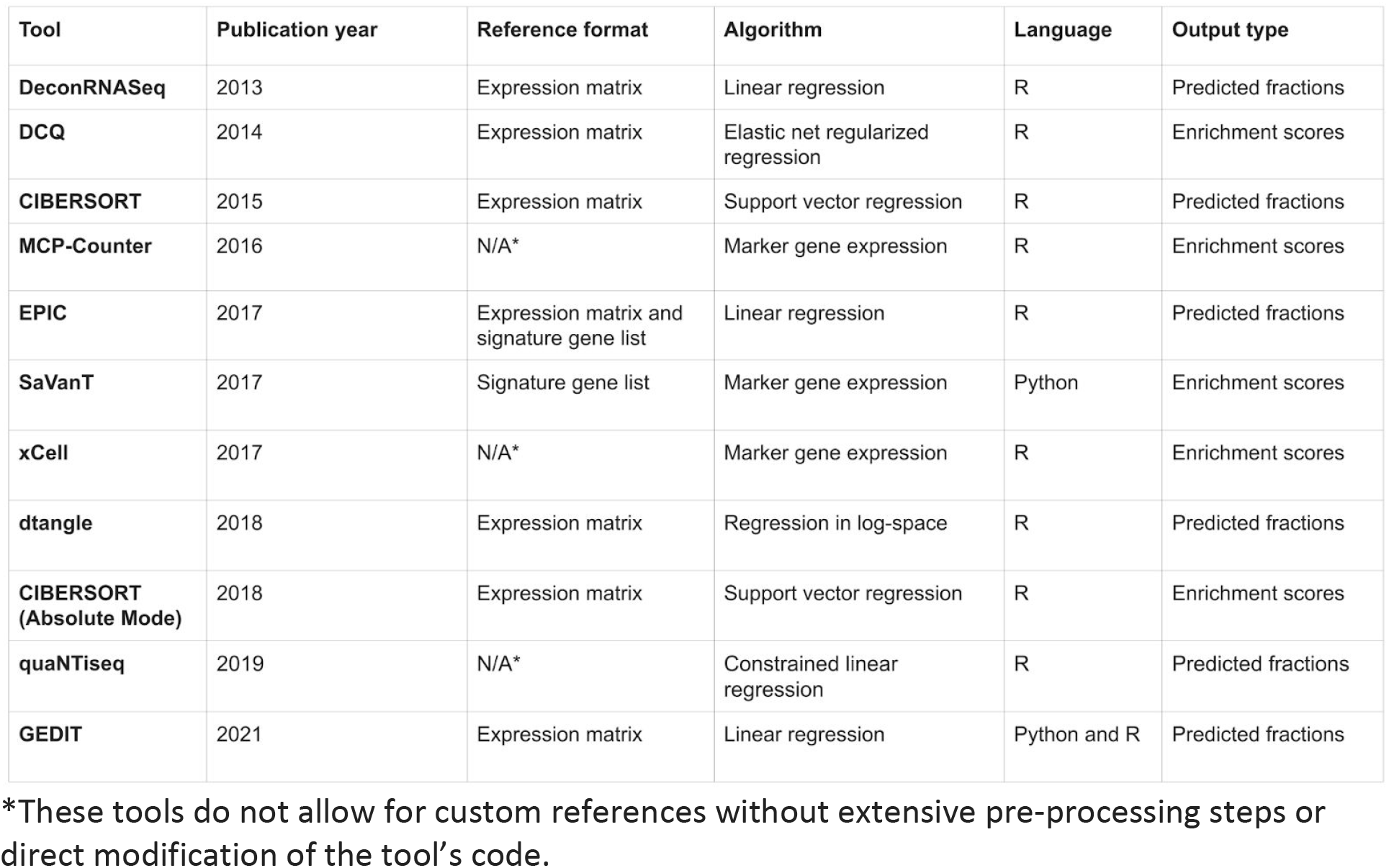
Summary of deconvolution tools evaluated by this benchmarking study, with details of inputs, algorithm, and publication. Included in the table are year of publication, format of required reference data, nature of the deconvolution algorithm and output values, and the language in which the tool is implemented. Tools are sorted by publication year. *These tools do not allow for custom references without extensive pre-processing steps or direct modification of the tool’s code.

Supplementary Table 1. Overview of the gold standard datasets. Each mixture comes from one of three platforms, and consists of 12-3728 samples of varying sets of cell types. Mixtures were either created *in silico*, mixed *in* vitro via titration, or taken directly from clinical patients. Underlying cell type fractions are either known naturally by nature of mixture generation, or evaluated by electrical impedance counting.

Supplementary Table 2. The set of reference matrices used in this study. These come from a variety of different platforms and publication sources, and each spans a different set of cells. Some references have been filtered to contain only a small set of signature genes ^24,31^. Others contain a larger set of tens of thousands of genes captured by the respective technology.

Supplementary Table 3. Area under curve values referenced in Figure 3.

Supplementary Table 4. Cell types present in each reference matrix used in this study.

Supplementary Figure 1. Single cell data used for generation of PBMC *in silico* mixtures. a) Cluster visualization using UMAP. b) Marker genes used for cluster identification.

Supplementary Figure 2. Cell type composition of 12 cell type mixtures created *in vitro*. Purified cell types were titrated and mixed in known proportions, followed by expression profiling on an Illumina HT12 BeadChip microarray

Supplementary Figure 3. Averaged absolute error values when eight deconvolution tools are executed using each of ten reference matrices.

Supplementary Figure 4. Scatter plots of predicted vs actual cell type fractions when eleven deconvolution tools are applied to *in vitro* cell mixtures.

Supplementary Figures 5. Scatter plots of predicted vs actual cell type fractions when eleven deconvolution tools are applied to *in silico* PBMC mixtures.

Supplementary Figures 6. Scatter plots of predicted vs actual cell type fractions when eleven deconvolution tools are applied to *in silico* Stromal mixtures

Supplementary Figure 7. Distribution of absolute error when deconvolution tools are applied to 3,728 samples from the Framingham cohort. True fractions of each cell type are evaluated by impedance-based electronic cell counting. For each tool requiring a reference, we selected the reference that produced the highest correlation between actual fractions and tool output.

