## Supplementary Materials for "Systematic evaluation of transcriptomics-based deconvolution methods and references using thousands of clinical samples"

Supplementary Table 1

| Dataset | Sequencing Platform | Cell Types | Number of Samples | Mixture Type | Cell Quantification Method |
|---|---|---|---|---|---|
| **Cell Mixtures** | Microarray | B, Mono, NK, Neutrophil, CD4 T, CD8 T | 12 | *In vitro* cell mixtures | Controlled cell mixing *in vitro* |
| **Simulated PBMC** | psuedo-bulk from scRNA-seq | B, Mono, NK, CD4 T, CD8 T | 200 | *In silico* simulated mixtures | Simulated cell mixing *in silico* |
| **Simulated Stromal Mixtures** | psuedo-bulk from scRNA-seq | B, CD4 T, CD8 T, Endothelial, Fibroblast, Macrophages, Mast cells | 100 | *In silico* simulated mixtures | Simulated cell mixing *in silico* |
| **Framingham Cohort Data** | Microarray | Neutrophils, Lymphocytes, Monocytes | 3728 | Human clinical samples | Electrical impedance counting |

Supplementary Table 2

| Reference | Platform | Assembly Publication | Cell Types | Number of Genes |
|---|---|---|---|---|
| **10x Immune** | Single Cell RNA-seq | Nadel *et. al.* 2021 | 9 Immune | 20,340 |
| **BLUEPRINT** | Bulk RNA-seq | Nadel *et. al.* 2021 | 8 Immune | 14,381 |
| **BlueCode** | Bulk RNA-Seq | Nadel *et. al.* 2021 | 35 stromal and immune | 13,300 |
| **Blood Circulating Immune Cells** | Bulk RNA-seq | Racle et al 2017 | 6 Immune | 49,900 |
| **Tumor Infiltrating Cells** | Bulk RNA-seq | Racle et al 2017 | 4 Immune<br>2 Stromal | 23,685 |
| **Human Primary Cell Atlas** | Affymetrix U133 Plus 2.0 | Mabbot *et. al.*, 2013 | 7 immune or 4 immune and 2 stromal (2 versions used) | 19,715 |
| **ImmunoStates** | Multi-Microarray | Vallania *et. al.*, 2018 | 20 immune | 317 |
| **LM22** | Affymetrix Microarray | Newman *et. al.*, 2017 | 22 immune | 547 |
| **Skin Signatures** | Multi-Microarray | Swindell *et. al.*, 2004 | 16 immune 5 stromal | 20,307 |

Supplementary Table 3

| | AbsCIBERSORT | CIBERSORT | DCQ | DeconRNASeq | EPIC | GEDIT | MCP-counter | SaVanT | dtangle | quanTIseq | xCell |
|---|---|---|---|---|---|---|---|---|---|---|---|
| **PBMC in silico mixtures** | 0.471 | 0.471 | 0.450 | 0.455 | 0.455 | 0.477 | 0.440 | 0.442 | 0.463 | 0.462 | 0.445 |
| **CellMixtures** | 0.468 | 0.467 | 0.455 | 0.453 | 0.464 | 0.462 | 0.448 | 0.458 | 0.459 | 0.459 | 0.456 |
| **Framingham Mixtures** | 0.444 | 0.444 | 0.383 | 0.462 | 0.439 | 0.466 | 0.440 | 0.385 | 0.370 | 0.444 | 0.401 |
| **Stromal in silico Mixtures** | 0.472 | 0.472 | 0.460 | 0.461 | 0.466 | 0.480 | 0.466 | 0.463 | 0.471 | 0.454 | 0.478 |

Supplementary Table 4

| Reference Source | Cell Types |
|---|---|
| 10x Immune | CD14 monocytes, CD19 B, CD4 Helper T, CD4 Memory T, CD4 Naive T, CD4 Regular, CD56 NK, CD8 Cytotoxic T, CD8 Naive T |
| BlueCode | Chondrocyte.articular B.cell.naive B.cell.memory Monocyte.CD14 NK.cell.CD56 Endothelial.large.blood.vessel Endothelial.microvascular.mammary.or.endometrial Endothelial.microvascular.non.reproductive Epithelial.cell.mammary Epithelial.renal.cortical Epithelial.respiratory Fibroblast.cardiac.ventricle Fibroblast.of.arm Fibroblast.of.lung Fibroblast.of.the.aortic.adventitia Fibroblast.or.papilla.dermal.cell Hair.follicular.keratinocyte Macrophage Mature.neutrophil Melanocyte.of.skin Myometrial.cell Osteoblast MSC.like.pluripotent.cell Myocyte.regular.cardiac Smooth.muscle.cell.aortic Smooth.muscle.cell.bronchialSmooth.muscle.cell.of.bladder Smooth.muscle.cell.coronary.artery Smooth.muscle.cell.pulmonary.artery Smooth.muscle.cell.of.trachea Smooth.muscle.cell.uterine T.Cell.CD4 T.cell.CD4.memory T.cell.CD8.memory |
| BluePrint | CD38.negative.naive.B.cell memory.B.cell CD14.positive..CD16.negative.classical.monocyte cytotoxic.CD56.dim.natural.killer.cell mature.neutrophil memory.CD4.positive.alpha.beta.T.cell CD4.positive..alpha.beta.T.cell memory.CD8.positive.alpha.beta.T.cell |
| EPIC - Blood Circulating Immune Cells | Bcells Monocytes NKcells Neutrophils CD4_Tcells CD8_Tcells |
| EPIC - Tumor Infiltrating Cells | BcellsCD4_Tcells CD8_Tcells Endothelial CAFs Macrophages |
| HPCA - Blood | B_cells Monocytes_CD14+CD16- Monocytes_CD16+CD14- NK_cells Neutrophils T_cells_(CD4+) T_cells_(CD8+) |
| HPCA - Stromal | B_cells T_cells_(CD4+) T_cells_(CD8+) Endothelial_cells Fibroblasts Macrophage |
| ImmunoStates | naive_B_cell memory_B_cell CD14_positive_monocyte CD16_positive_monocyte CD56dim_natural_killer_cell CD56bright_natural_killer_cell neutrophil CD4_positive_alpha_beta_T_cell CD8_positive_alpha_beta_T_cell macrophage_m0 macrophage_m1 macrophage_m2 MAST_cell basophil eosinophil gamma_delta_T_cell hematopoietic_progenitor myeloid_dendritic_cell plasma_cell plasmacytoid_dendritic_cell |
| LM22 | B cells naive B cells memory Monocytes NK cells resting NK cells activated Neutrophils T cells CD4 naive T cells CD4 memory resting T cells CD4 memory activated T cells CD8 Macrophages M0 Macrophages M1 Macrophages M2 Mast cells resting Mast cells activated Plasma cells T cells follicular helper T cells regulatory (Tregs) T cells gamma delta Dendritic cells resting Dendritic cells activated Eosinophils |
| Skin Signatures | B_cells Blood_CD34._cells Dendritic_cells Eosinophils Fibroblasts Keratinocytes Lymphatic_endothelial_cells Macrophages Monocytes NK_cells Neutrophils Plasma_cells_CD138. Platelets Reticulocytes Subcutaneous_Adipose T_cells_CD3. T_cells_CD4. T_cells_CD8. T_cells_Gamma_Delta T_cells_Regulatory Vascular_Endothelial_Cells |

Supplementary Figure 1

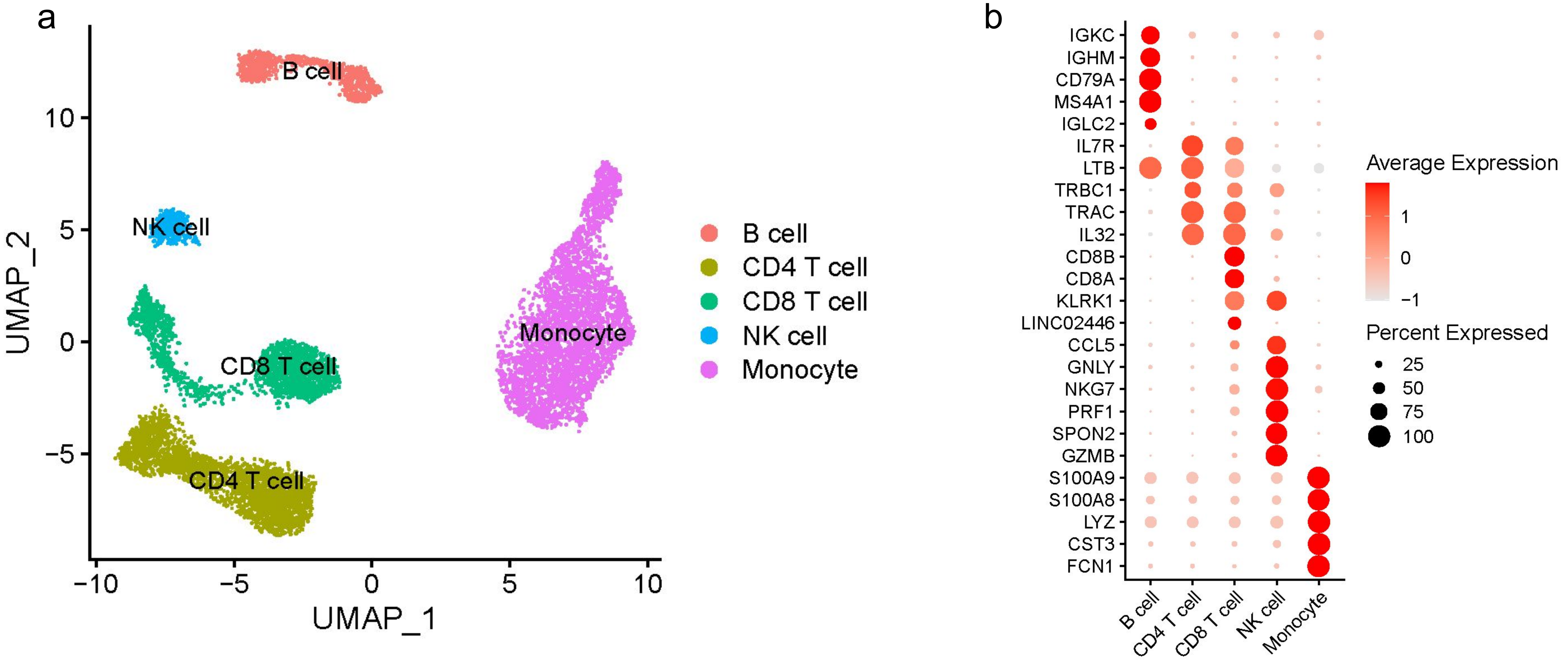

Supplementary Figure 2

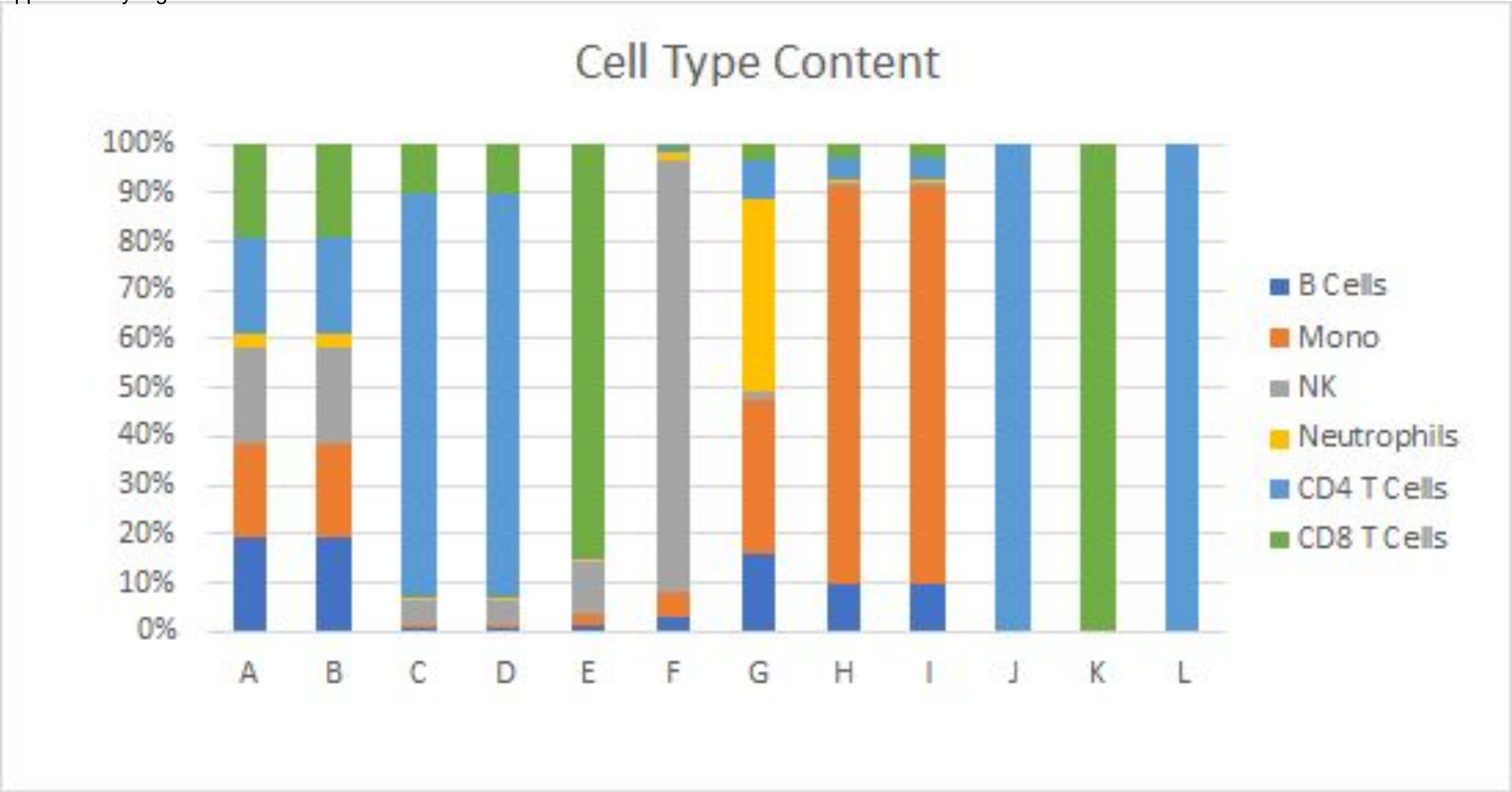

Supplementary Figure 3

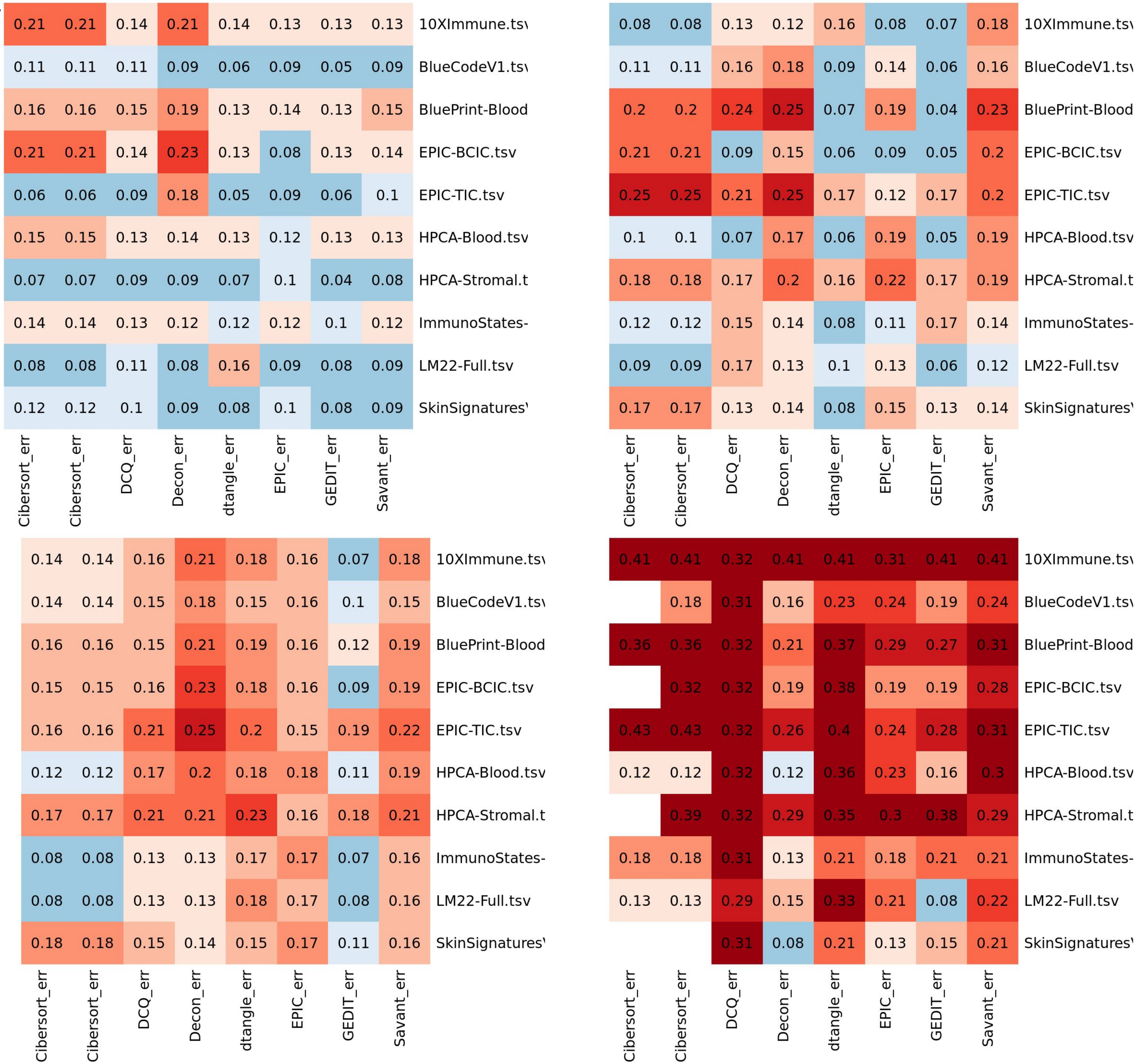

Supplementary Figure 4

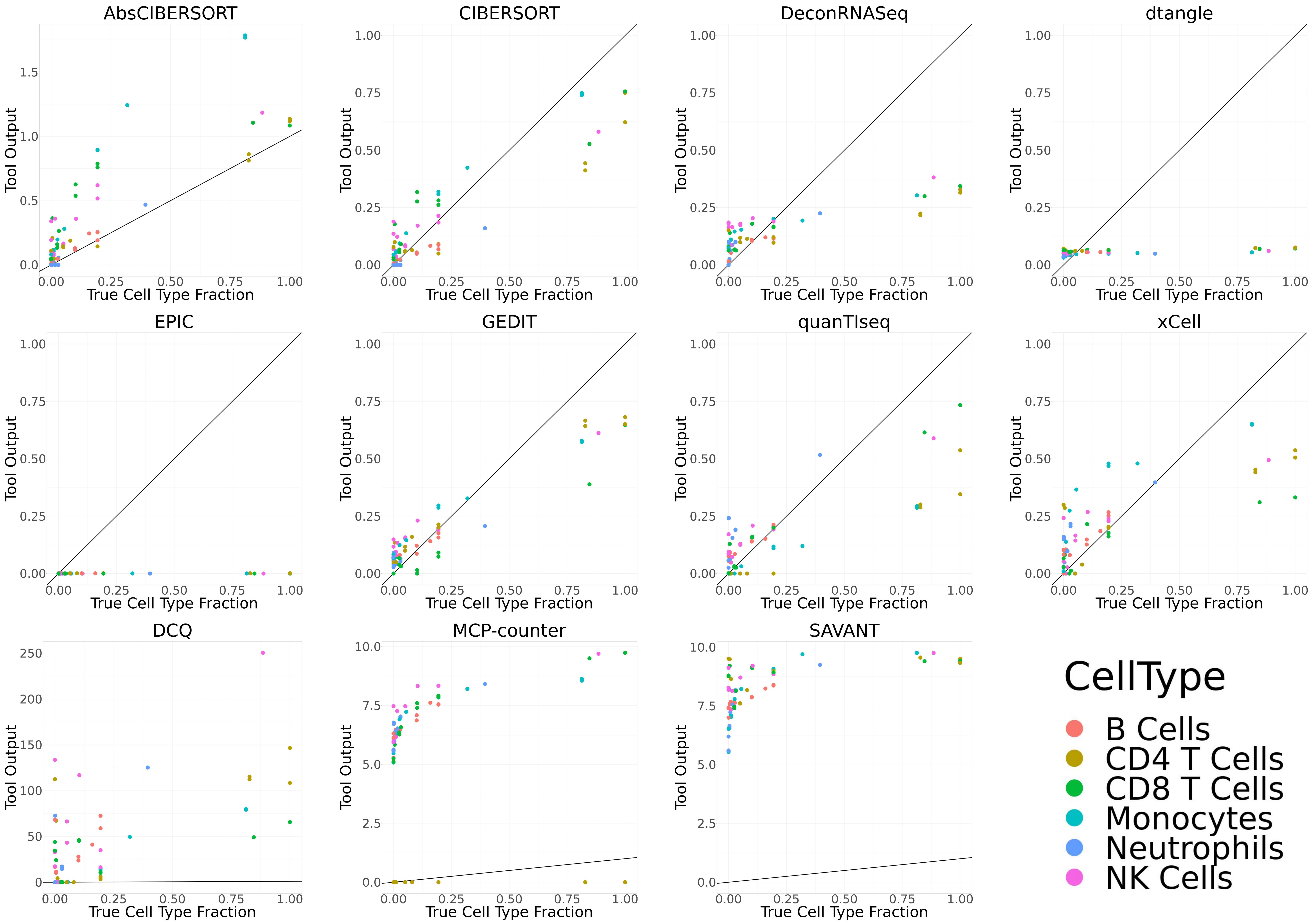

Supplementary Figure 5

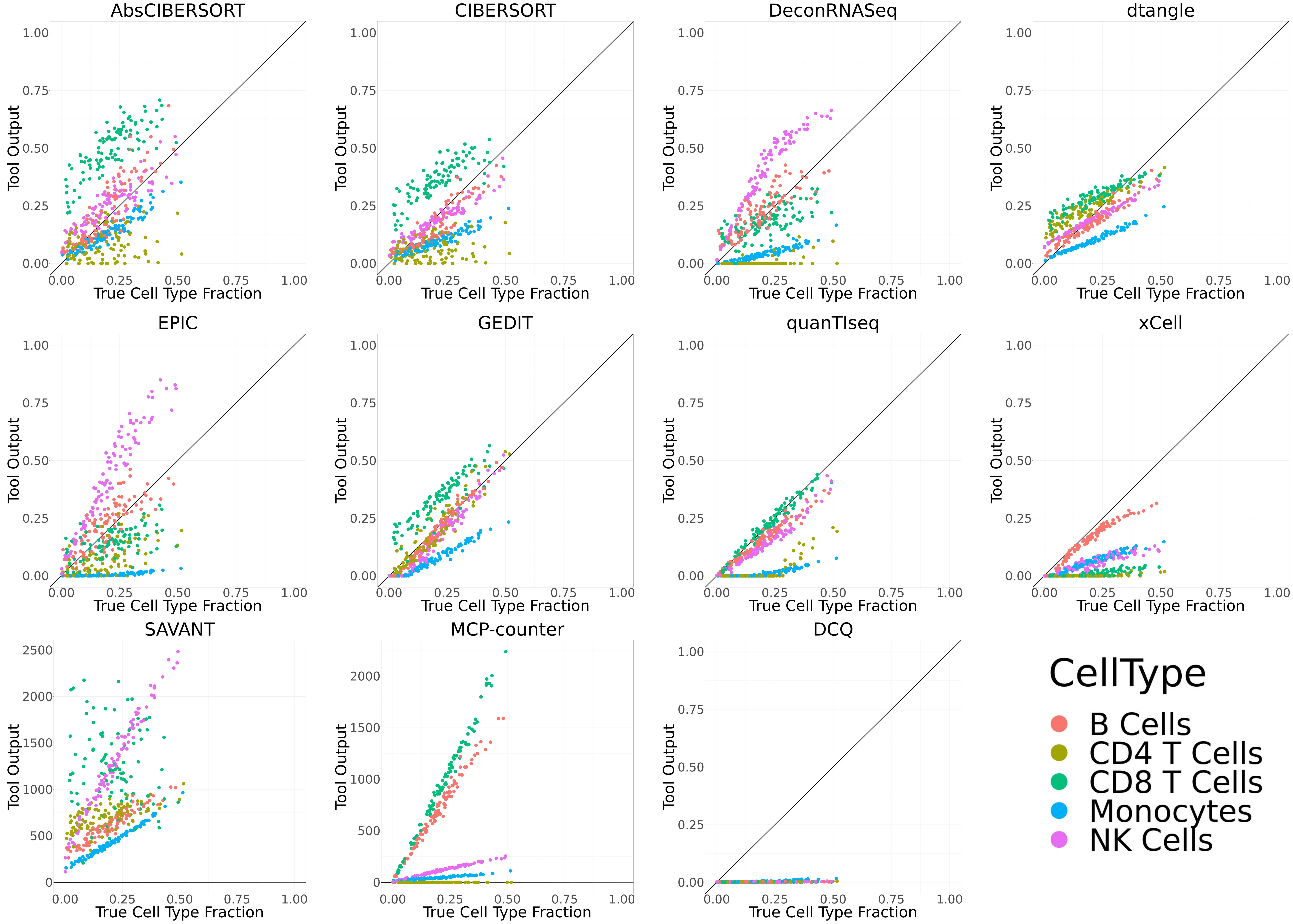

Supplementary Figure 6

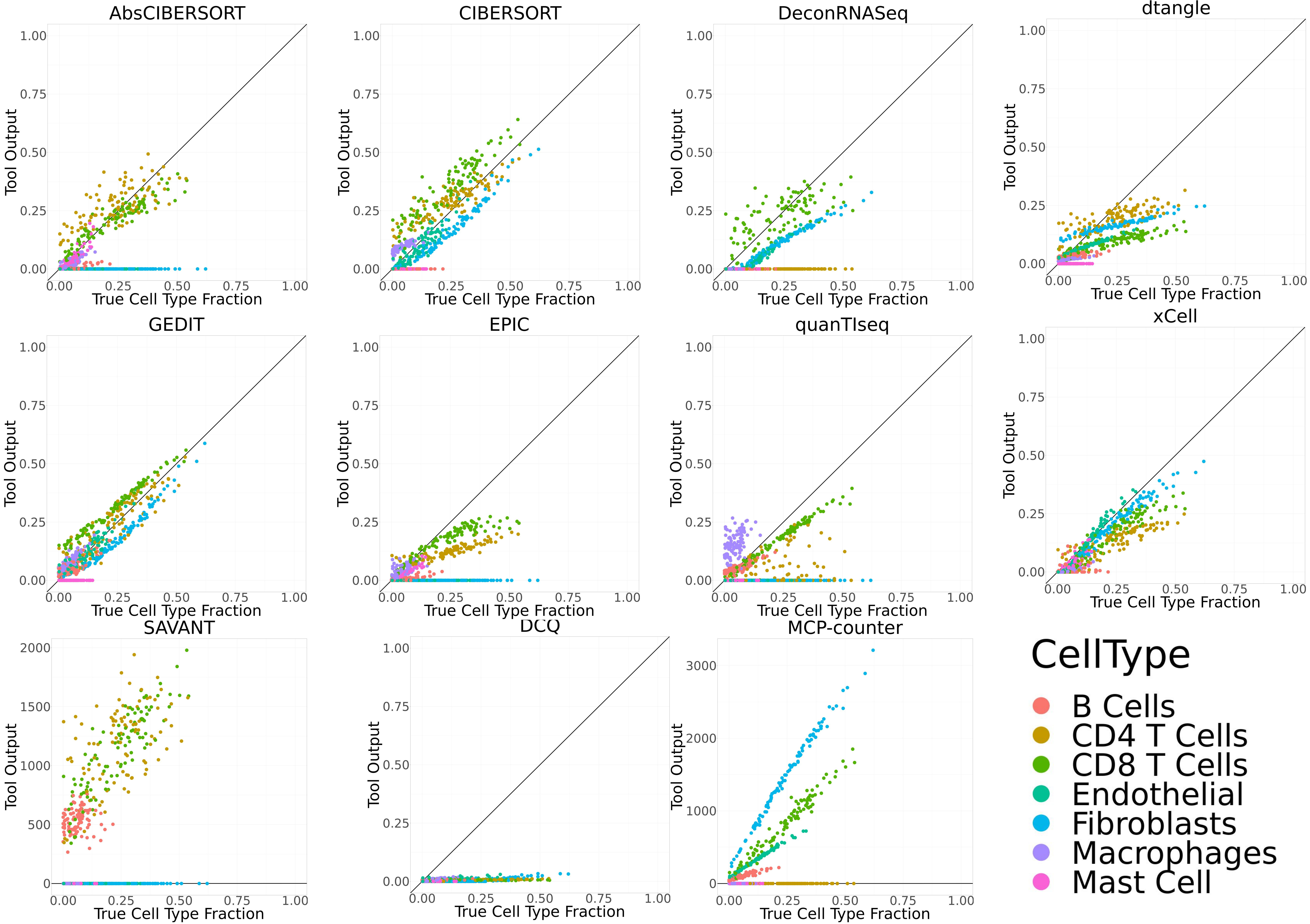

Supplementary Figure 7

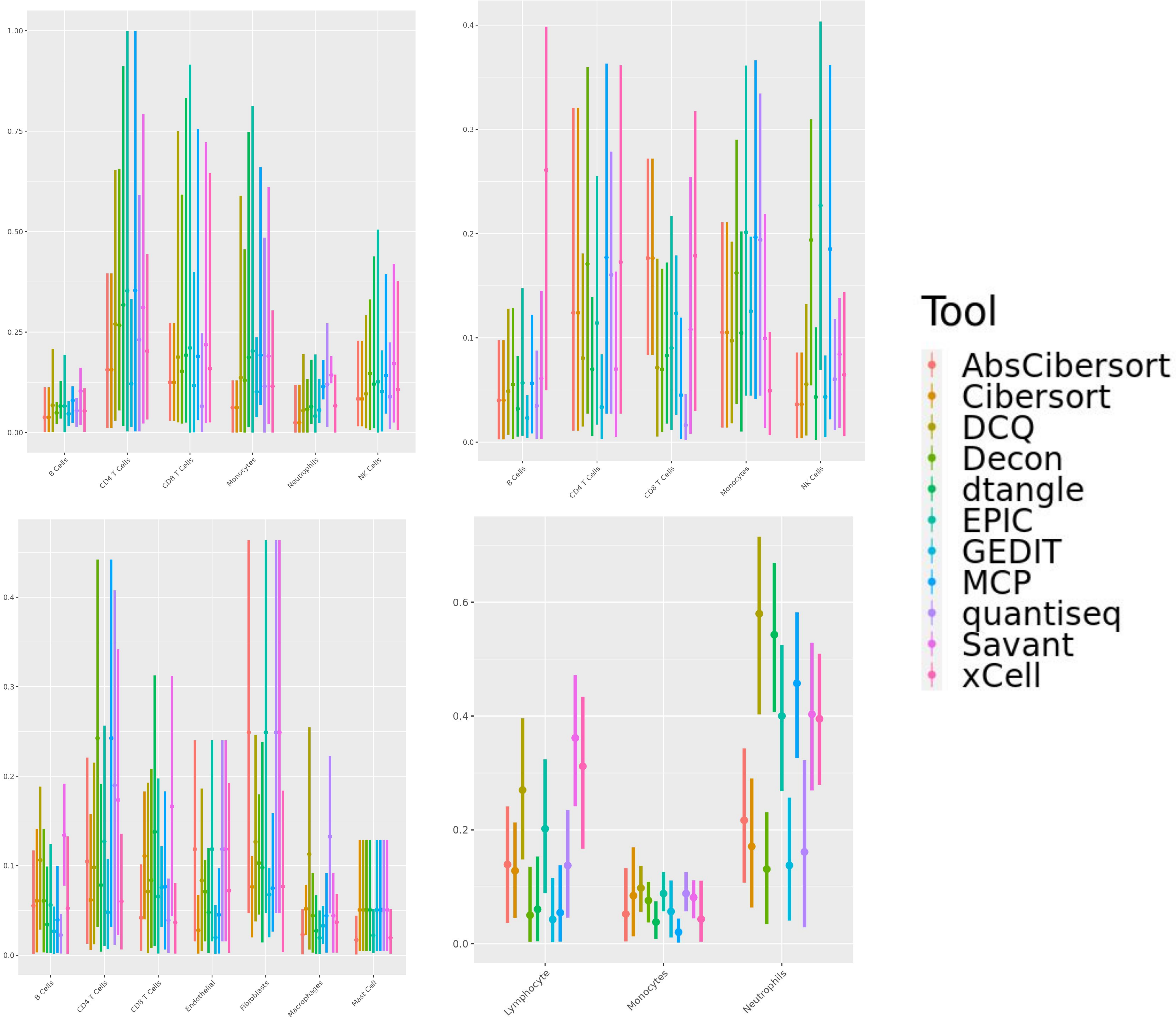

Supplementary Table 1. Overview of the gold standard datasets. Each mixture comes from one of three platforms, and consists of 12-3728 samples of varying sets of cell types. Mixtures were either created *in silico*, mixed *in* vitro via titration, or taken directly from clinical patients. Underlying cell type fractions are either known naturally by nature of mixture generation, or evaluated by electrical impedance counting.

Supplementary Table 2. The set of reference matrices used in this study. These come from a variety of different platforms and publication sources, and each spans a different set of cells. Some references have been filtered to contain only a small set of signature genes [24,31]. Others contain a larger set of tens of thousands of genes captured by the respective technology.

Supplementary Table 3. Area under curve values referenced in Figure 3.

Supplementary Table 4. Cell types present in each reference matrix used in this study.

Supplementary Figure 1. Single cell data used for generation of PBMC *in silico* mixtures. a) Cluster visualization using UMAP. b) Marker genes used for cluster identification.

Supplementary Figure 2. Cell type composition of 12 cell type mixtures created *in vitro*. Purified cell types were titrated and mixed in known proportions, followed by expression profiling on an Illumina HT12 BeadChip microarray

Supplementary Figure 3. Averaged absolute error values when eight deconvolution tools are executed using each of ten reference matrices.

Supplementary Figures 4. Scatter plots of predicted vs actual cell type fractions when eleven deconvolution tools are applied to *in vitro* cell mixtures.

Supplementary Figures 5. Scatter plots of predicted vs actual cell type fractions when eleven deconvolution tools are applied to *in silico* PBMC mixtures.

Supplementary Figures 6. Scatter plots of predicted vs actual cell type fractions when eleven deconvolution tools are applied to *in silico* Stromal mixtures

Supplementary Figure 7. Distribution of absolute error when deconvolution tools are applied to 3,728 samples from the Framingham cohort. True fractions of each cell type are evaluated by impedance-based electronic cell counting. For each tool requiring a reference, we selected the reference that produced the highest correlation between actual fractions and tool output.
